# Optimization of chickpea irrigation in a semi-arid climate based on morpho-physiological parameters

**DOI:** 10.1101/2023.02.12.528176

**Authors:** Asaf Avneri, Zvi Peleg, David Bonfil, Roy Sadeh, Omer Perach, Ittai Herrmann, Shahal Abbo, Ran N. Lati

## Abstract

While the world population is steadily growing, the demand for plant-based protein in general, and chickpea in particular, is rising. Heatwaves and terminal drought are the main environmental constraints on chickpea production worldwide. Thus, developing better irrigation management for the chickpea agro-system can promote higher and more sustainable yields. Supplemental irrigation at the right timing and dose can increase yield dramatically. Here, we studied the response of a modern Kabuli chickpea cultivar to supplemental irrigation during the critical pod-filling period over three growing seasons (2019-2021) in northern Negev, Israel, under semi-arid conditions. Six irrigation treatments were applied based on irrigation factors of 0, 0.5, 0.7, 1.0, 1.2, and 1.4 of crop evapotranspiration (ET_0_) as measured by an on-site meteorological station. Morpho-physiological parameters and above-ground biomass accumulation were monitored throughout the cropping seasons, and the final grain yield was determined at maturation. Irrigation onset was determined based on plants’ leaf water potential (Ψ^LWP^ > 15 bar) in the field. Our results indicate that optimal water status (as reflected by pressure chamber values) was 12-14 bar during the irrigation period. Irrigation according to evapotranspiration (ET_0_) with an irrigation factor of 1.2 resulted in the highest grain yields over the three years. To ensure optimal water supply during the reproductive phase compatible with the crop water requirements, maintaining a 25 mm node length above the last fully developed pod and a 90 mm distance between the last fully developed pod to the stem apex is recommended. In conclusion, irrigation onset when the crop is already at mild drought stress, followed by sufficient irrigation while following the indicated morphology and water potential values, may help farmers optimize irrigation and maximize chickpea crop production.

**Highlights:**

- Chickpea irrigation is optimized based on meteorological data (ETc) and morphological traits.
- Morphological- not only physiological- traits can capture the dynamics of crop water status.
- Irrigation onset should be at mild drought stress (>15 bar) during the reproductive stage.
- Extending the reproductive phase is essential for grain yield improvement under semi-arid conditions.
- IF=1.2 provides the most productive irrigation regime for chickpea in Mediterranean conditions.

## 1. Introduction

World agriculture faces the challenge of feeding the growing global population under diminishing resources and changing climate, as reflected in the increasing frequency and intensity of extreme climatic phenomena such as droughts and heatwaves (Sade and Peleg, 2020). In the eastern Mediterranean, a significant reduction in the number of seasonal rain events is accompanied by an increase in rain intensity (per event) (Drori et al., 2021), resulting in lower soil water availability for rainfed crops.

Irrigated farmland is estimated at 20-25% of the world’s cultivated land, producing ∼40% of agricultural yield (Puy et al., 2021). Timely irrigation may enhance crop yield compared with rainfed farming (Mueller et al., 2012), but the availability and timing of water resources limit farmers’ options. One way to improve global food security is to enlarge the area of irrigated staple crops, making them less reliant on precipitation and with higher potential yield. One such crop is the chickpea (*Cicer arietinum* L.), an important source of dietary protein (van der Maesen, 2007), which is getting more attention as a healthy and functional food. In parallel, there is an increasing demand for plant-based proteins (like chickpea) to replace animal proteins (Bar-El Dadon et al., 2017). This calls for agronomic research to develop advanced agro-techniques to promote yield (Singh et al., 2016).

The global chickpea production is about 14.2 million tons, harvested from ca. 13.7 million hectares (FAO stat 2021), mostly grown on marginal lands in semi-arid regions. Under traditional farming, chickpea is a post-rainy season crop (Kostrinski, 1974), a strategy adopted in antiquity to avoid the damage from ascochyta blight, a disease caused by the pathogen *Didymella rabiei* which spreads by raindrops (Abbo et al., 2003). Consequently, most chickpea cropping relies on residual soil moisture without adequate irrigation infrastructure and water resources. The gap between the high yield potential of chickpea (over 4000 kg/ha; Singh, 1990) and the average global is considered a combined result of biotic and abiotic stresses (Singh and Jana, 1993), with water availability during the reproductive phase as the most critical yield-limiting factor (Awasthi et al., 2014). Efficient and cost-effective irrigation is a key strategy to help close the gaps between yield potential and actual yields in semi-arid agro-systems.

Estimation of crop water requirement by calculating evapotranspiration (ET) from meteorological data relies on the reference ET (ET_0_) (Doorenbos, 1977). Crop evapotranspiration (ET_0_) can be derived from meteorological data and expressed in the Penman-Monteith equation as the ET from a well-watered grass surface with fixed crop height, albedo, and surface resistance. Standard crop evapotranspiration (ET_c_) is defined as evapotranspiration under standard conditions (ETc) and refer to crop grown in a large field with optimal soil, water, and agronomical management that have a full production potential under the given climatic conditions (Allen et al., 1998). Many factors affect the actual field evapotranspiration, including (but not limited to) the crop species, crop cultivar, leaf area index (LAI), developmental stage, canopy height, crop roughness, a reflection of solar radiation, and ground cover. Consequently, under identical environmental conditions, different ET levels are expected for diverse crops or growing stages and should be considered when evaluating the evapotranspiration from crop fields. To bridge the gap between the reference evapotranspiration (ET_0_) and the crop evapotranspiration (ET_c_), a crop coefficient factor (Kc) is integrated. There are two crop coefficient (Kc) models, single or dual (Allen et al., 1998). Since during our irrigation period, the fraction of soil evaporation is low due to the full soil coverage by plants and the lack of daily water balance measurements of the evaporative soil surface layer, we choose to use the single crop coefficient model. The Kc is defined as the ratio of ET_c_/ET_0_ (Kc = ET_c_ / ET_0_ or ET_C_ = ET_0_ × Kc) (Allen et al., 1998). While the ET_0_ (based on the Penman-Monteith equation) takes into account the climatic parameters, the Kc represents the crop characteristics such as crop height, albedo (LAI effect), canopy resistance (leaf area, leaf anatomy, stomatal characteristics), (Allen et al., 1998). Thus, The Kc factor for a given crop changes throughout the crop life cycle by the crop growth stages and can correctly estimate the crop evapotranspiration (ETc). While the Kc matches climatic conditions and actual evapotranspiration in the field, here we examined the crop response to increasing irrigation factor (IF) of the cumulative ET_0_ (computed from the meteorological station) to promote the highest grain yield.

Due to its indeterminate growth habit, yield accumulation in chickpea depends upon simultaneous vegetative and reproductive development since flowering commencement. Therefore, supplemental irrigation is essential for extending the reproductive period and supporting higher grain and total biomass yields. Yet, it is still unclear when to irrigate and what water cubature should be applied at each irrigation to ensure maximum grain yield. In other summer crops, such as cotton (*Gossypium hirsutum*), two morphological traits (number of nodes above the last white flower and the distance of the white flower from the plant apex) are used to monitor the crop development and make real-time decisions concerning irrigation, and management (Bourland et al., 1992; Waddle, 1982). Previously, Bosak et al. (1996) suggested morpho-developmental criteria as a tool to monitor the water status and adjust chickpea irrigation. These authors mentioned that the chickpea’s Last Fully Developed Pod (LFDP) distance to the plant apex correlates with the plant water status. Yet, their research lacked actual evidence for that relationship, and the yield factor was overlocked.

We hypothesized that a gradient of irrigation treatments would result in a range of yield levels that would facilitate the determination of the agro-economic optima. Our underlying assumption was that crop morpho-physiological parameters would change alongside final grain yield with water availability treatments and their interaction with the seasonal weather conditions. Accordingly, our primary goal was to establish a protocol for optimizing chickpea irrigation, striving for maximum yield. In parallel, we calculated the water productivity (WP) as grain yield divided by the amount of available water (precipitation and irrigation) of the different irrigation factor treatments (Fernández et al., 2020; Pereira et al., 2012). We seek to identify morphological indices that may help monitor the crop water status and to develop chickpea irrigation guidelines for a semi-arid Mediterranean climate.

## 2. Materials and methods

### 2.1 Plant materials and growing conditions

The Israeli cultivar “Zehavit” (Kabuli type, Hazera LTD), with an average weight of 370 mg/seed, was used for the field experiments because it is adapted to a semi-arid climate and is the most common in Israel.

A three-year study was conducted during the 2019-2021 seasons at Gilat Research Center, ARO, Israel (34°39’ E; 31°20’ N), located 160 meters above sea level (Fig. 1). This area is characterized by mild winters with an average annual precipitation of 230 mm (Bonfil et al., 1999) while the spring and summer seasons are warm and dry (no rains at all). The soil at the location is sandy loam loess–Calcic Xerosol (Dan et al., 1982), with a pH of 7.9-8.3 and a bulk density of 1.45 g cm^-3^. Daily meteorological data were collected from the Gilat meteorological station, located within 100-1500 meters of the experiment fields, provided by the Israeli Ministry of Agriculture (http://www.meteo.moag.gov.il). The previous crop was bread wheat (*Triticum aestivum*) for all three experiments. As wheat utilizes all available water (Bonfil et al., 1999), we assumed that no residual water in the soil from the previous season was available to the chickpea plants (especially in soil layers populated by chickpea roots).

**Figure 1.**
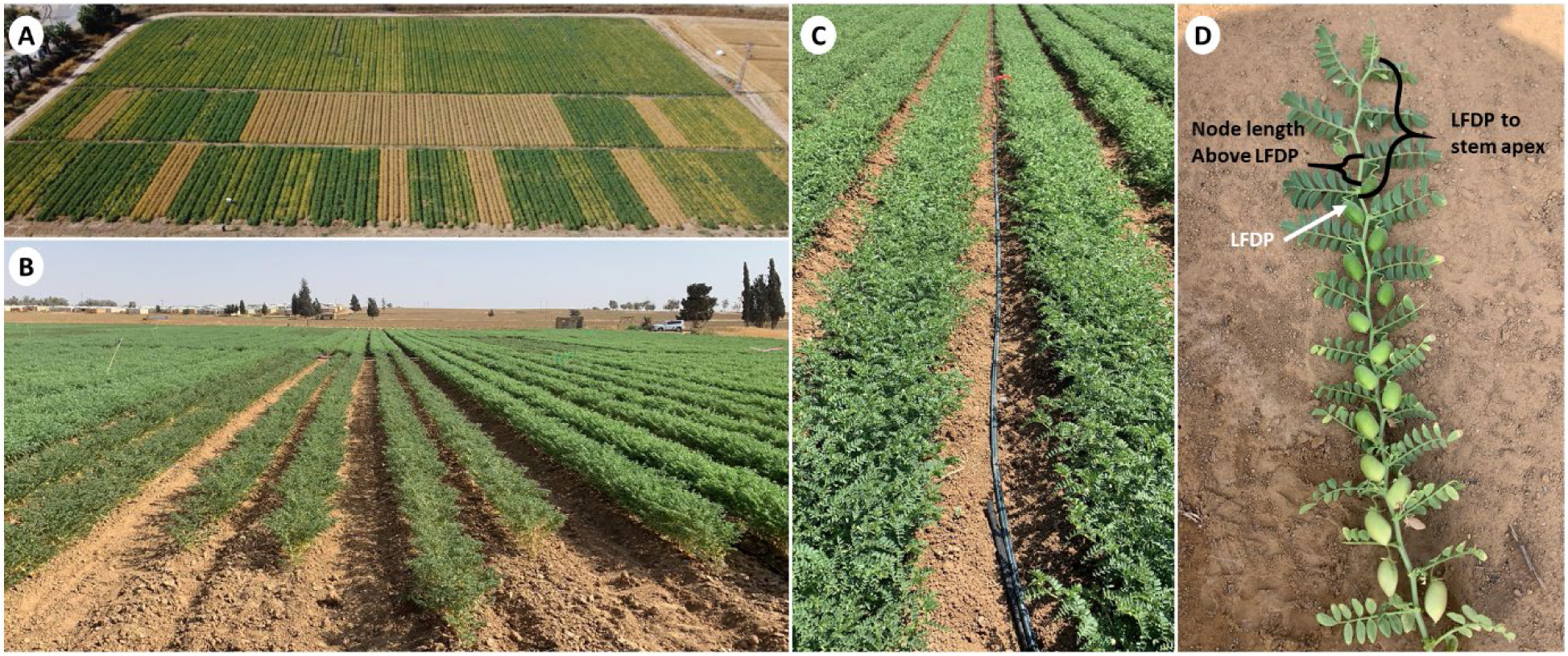
**A**) Aerial image of the field experiment (Gilat 2020). **B**) Overview of the field plots (Gilat 2021). **C**) Image of the dripping line located between rows. **D)** Image of a representative main branch. The last fully developed pod (LFDP), node length above LFDP, and distance from LFDP to stem apex are indicated.

### 2.2 Experimental design

The experiments were conducted in a complete randomized block design with six replicates. Each plot was 5.79 meters in width (three 1.93 meters in width beds) and 25 meters in length. Sowing was held with a row planter in 15 seeds/meter-row density on two rows/bed with a 75 cm distance between them. The experiments were initiated on the last week of January (25, 30, and 22 for 2019, 2020, and 2021, respectively), and seeds emerged at the beginning of February (03, 16, and 02 for 2019, 2020, and 2021, respectively). Between seed emergence and the reproductive stage, rain and supplemental sprinkler uniform irrigations were applied to ensure proper development before the onset of the differential drip irrigation. Differential drip-irrigation was applied weekly with six irrigation treatments that we defined by irrigation factors (IF) of 0, 0.5, 0.7, 1, 1.2, and 1.4 of the cumulative ET_0_ (computed from the meteorological station) since the last irrigation (Fig. S2) (in 2019 season, we did not have an IF=0 treatment). The differential irrigation was initiated only after leaf water potential (pressure chamber values) reached values above Ψ^LWP^ = 15 Bar. We selected this threshold based on agronomical knowledge in Israel. While facing heat waves (high temperatures and high evapotranspiration), irrigation was applied twice a week. For each irrigation factor treatment, we used a drip line with different dripping spaces (0.7, 0.5, 0.5, 0.3, and 0.5 meters between drippers, for IF 0.5, 0.7, 1, 1.2, and 1.4, respectively) and flow rate [0.7, 0.7, 1.0, 0.7, and 0.7 (two dripping lines/bed) liter per hour, for IF 0.5, 0.7, 1, 1.2, 1.4 respectively]. With different combinations, we managed to irrigate each treatment simultaneously with different cubature but for the same duration of irrigation. The first irrigation (see below for timing determining) amounts were calculated by measuring evapotranspiration a week before starting irrigation. Irrigation cubature was calculated according to evapotranspiration (ET_0_ values) from the previous irrigation until the date of application-multiplied by the specific IF for every irrigation treatment.

### 2.3 Morpho-physiological characterization

Leaf water potential (Ψ^LWP^) was measured weekly with a pressure chamber (Model 600D, PMS instrument company, OR, USA). One main stem from every plot was cut at the fifth or fourth node from the apex during the hottest part of the day (12:00-15:00) and quickly (<1 min) inserted into the chamber. The distance of the last fully developed pod (LFDP) from stem apices and the node length above LFDP were measured once a week following the appearance of the first pods (2019 and 2020 seasons) or after the first irrigation (2021 season) until full maturity of each treatment (Fig. 1D). Plant samples of 0.5-meter row were harvested weekly at ground level, dried for 48 hours at 70°C, and weighed to obtain total dry matter (Tot. DM). Later, the samples were threshed using a laboratory threshing machine (LD 350, WinterSteiger, Reid, Austria), and reproductive DM (Rep. DM) was weighed. Vegetative DM was calculated as the difference between total DM to reproductive DM. At full maturity, plots were harvested by a conventional grain harvester to obtain the grain yield (GY). The harvest index (HI) was calculated as the ratio between Rep. DM and Tot. DM. Water productivity (WP) was calculated as the ratio between grain yield and water used (sum of irrigation and rainfall) for each irrigation treatment.

### 2.4 Statistical analyses and data visualization

The JMP^®^ ver. 16 statistical package (SAS Institute, Cary, NC, USA) was used for statistical analyses unless specified otherwise. A factorial model was employed for the analysis of variance (ANOVA), with irrigation regimes as a fixed effect and years considered random effects. Descriptive statistics are graphically presented in the box plots: median value (horizontal short line), quartile range (25 and 75%), and data range (vertical long line).

## 3. Results

### 3.1 Climatic conditions and irrigation pattern throughout the three growing seasons

The seasonal pattern of daily minimum and maximum temperatures during the vegetative stage before the application of differential drip irrigation shows that the 2019 season was characterized by a relatively mild spring (range 9.4-21.8°C) which allowed an onset of irrigation at 79 days after emergence (DAE) compared with 75 and 62 DAE in 2020 (range 10.1-23.4°C) and 2021 (range 8.7-21.8°C), respectively. This earlier irrigation onset in the 2021 season corresponded with the lower water availability; precipitation (140.7 mm), and sprinkler irrigation (60 mm). The evapotranspiration (ET_0_) ranged from 263.1, 250.2, and 188.7 mm for the 2019, 2020, and 2021 growing seasons, respectively (Fig. S1). In 2019, temperatures and evapotranspiration data showed mild conditions during the vegetative stage before applying differential drip irrigation. This allowed the gradual development of plants without extreme stress episodes. As the season progressed, the crop faced several heat waves (Fig. S1A). The 2020 seasonal temperature and ET_0_ pattern show an extreme 8-day ongoing heat wave (accompanied by high ET_0_ values) less than two weeks after commencing irrigation (88 DAE). These extreme climatic conditions resulted in the restricted canopy, pod development, and grain set (Fig. S1B). In the 2021 season, the crop experienced several heat waves, two of which occurred before the onset of irrigation (62 DAE), followed by several events during the irrigation season until maturity (Fig. S1C). The average minimum temperatures were similar across the three growing seasons (11.3°C), while the average maximum temperatures ranged from 25.7 (2019) to 26.1°C (2020).

During the 2019 and 2021 growing seasons, the irrigation treatment of IF=1 [302.0 and 389.5 mm (liters m^-2^), respectively] was very close to measured reference evapotranspiration (ET_0_) during the irrigation period (310.9 and 373.2 mm, respectively; Fig. S2). In the 2020 season, due to extreme heat waves (i.e., days with maximum temperature > 35°C, ET_0_ > 6.5 mm/day), we had to apply a higher water cubature (277.5 mm irrigation compared with ET_0_ of 228.7; Fig. S1), trying to keep plants vital and functioning. The hostile evapotranspiration values during 2020 affected the plants’ growth and development, resulting in a 28 days irrigation period, while the irrigation periods in the 2019 and 2021 seasons were 42 and 54 days, respectively (Table S1).

### 3.2 Physiological characterization

In general, analysis of variance (ANOVA) resulted in a significant effect of the irrigation treatment and year on productivity (rep. DM, total DM, and GY), physiological (leaf water potential and water-use efficiency), and morphological (distance from apex to last inflated pod and length of node above the last inflated pod) parameters, except for harvest index (Table 1). A significant irrigation × year interaction effect was found for GY, Ψ^LWP^, WP, and the distance from apex to last inflated pod and length of node above the last inflated pod (Table 1). Yearly analyses of variance to test the effect of differential irrigation treatment showed a significant effect for all parameters except harvest index, which was significant only during the 2021 season (Table S2).

**Table 1.** Analysis of variance (ANOVA) for total dry matter (Tot. DM), reproductive DM (Rep. DM), grain yield (GY), harvest index (HI), instantaneous leaf water potential at median irrigation period (Ψ^LWP^), water productivity (WP), the distance of the last fully developed pod (LFDP) from plant-apex and node length above LFDP at median irrigation period, under different irrigation factors (0.5, 0.7, 1, 1.2, and 1.4) over three experimental seasons (2019, 2020, and 2021).

| Source | d.f. | Sum of square |  |  |  |  |  |  |  |
| --- | --- | --- | --- | --- | --- | --- | --- | --- | --- |
| | | Tot. DM | Rep. DM | GY | HI | $\Psi^{LWP}$ | WP | LFDP-stem apex | Node above LFDP |
| Irrigation (I) | 4 | 363079356<br>0.0001 | 110228322<br>0.0001 | 51482379<br>0.0001 | 0.017021<br>0.1236 | 525.51<br>0.0001 | 0.0622889<br>0.0127 | 71996.1<br>0.0001 | 1508.18<br>0.0001 |
| Year (Y) | 2 | 67979614<br>0.0001 | 29865018<br>0.0001 | 8731135<br>0.0001 | 0.023848<br>0.0073 | 495.57<br>0.0001 | 0.6771756<br>0.0001 | 54215.6<br>0.0001 | 1729.8<br>0.0001 |
| I×Y | 8 | 20992203<br>0.2579 | 6673436<br>0.2814 | 6091376<br>0.0001 | 0.009<br>0.8562 | 63.76<br>0.0001 | 0.130<br>0.0015 | 17012.2<br>0.0001 | 508.42<br>0.0001 |
| Error | 75 | 151662642 | 49959031 | 7926023 | 0.170 | 80.26 | 0.342 | 7141.7 | 278.5 |

d.f., degree of freedom.

During the 2020 and 2021 seasons, instantaneous leaf water potential values at the onset of irrigation matched our pre-planned value (pressure-chamber values above 15 Bar). This value represents levels of water stress in farmers’ fields that may impair the continuation of reproductive growth. The leaf water potential responded coherently to the irrigation treatments: high pressure-chamber values under deficit irrigation treatments compared with low values under optimal (well-irrigated) treatments. Under irrigation treatments of IF>1, the pressure-chamber values during the irrigation period ranged around 12-13.5 bar, attesting to the plants’ improved water status (Fig. 2).

**Figure 2.**
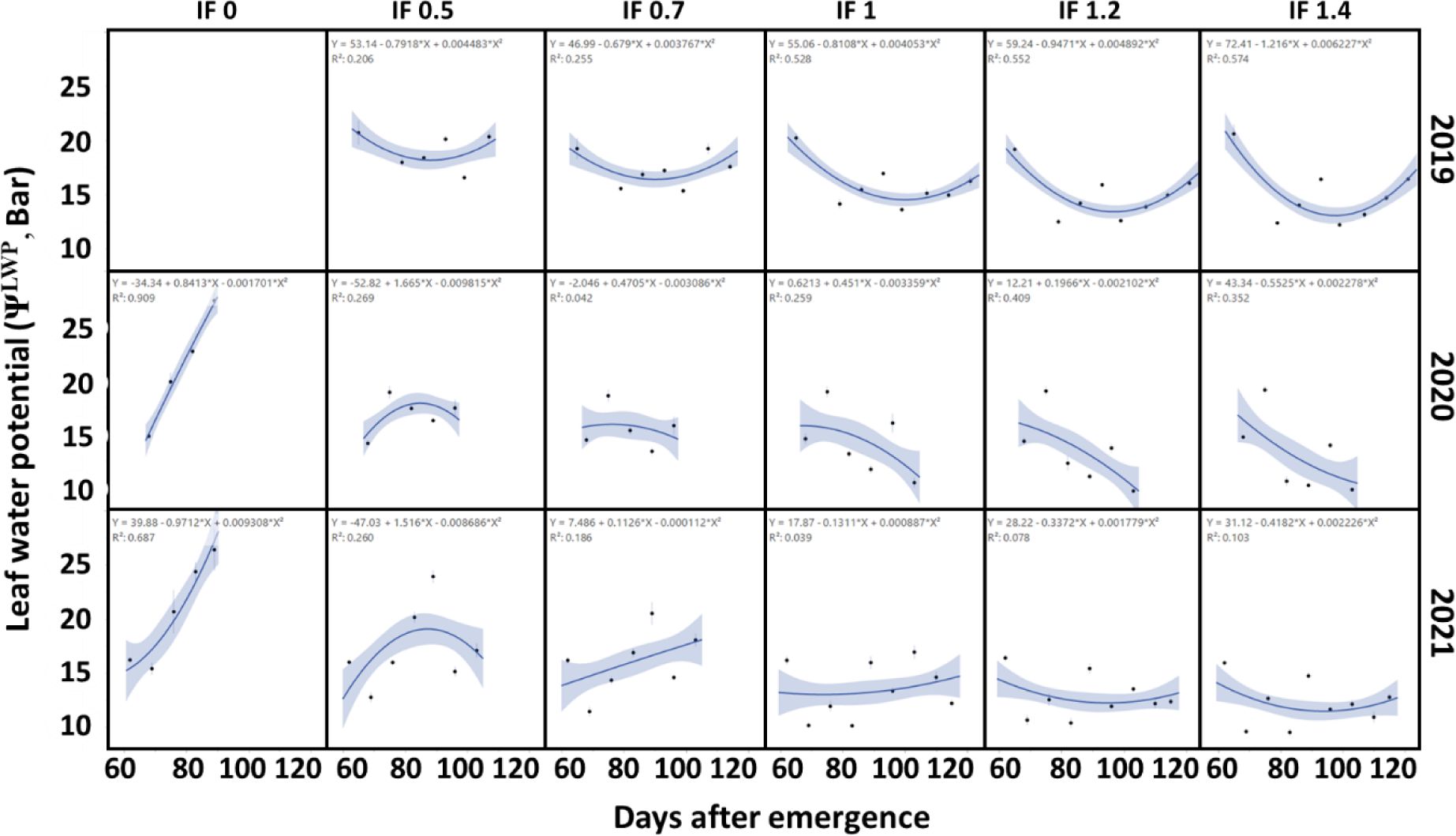
Instantaneous leaf water potential (pressure chamber values, Bar) along the irrigation period for each irrigation factor (IF= 0, 0.5, 0.7, 1, 1.2, and 1.4) over three experimental seasons (2019, 2020, and 2021). The curves represent the mean within the water treatment and year (*n*=6). Shading indicates the standard error.

### 3.2 Morphol-development parameters

The dynamics of the node length above the youngest fully inflated pod are presented in Figure 3A. For this morphometric parameter, the water treatments resulted in two responsiveness groups, one with IF≥1, which showed increased values following irrigation throughout the irrigation period. The second response group comprised the irrigation treatments with IF<1 and showed continuous growth decay.

**Figure 3.**
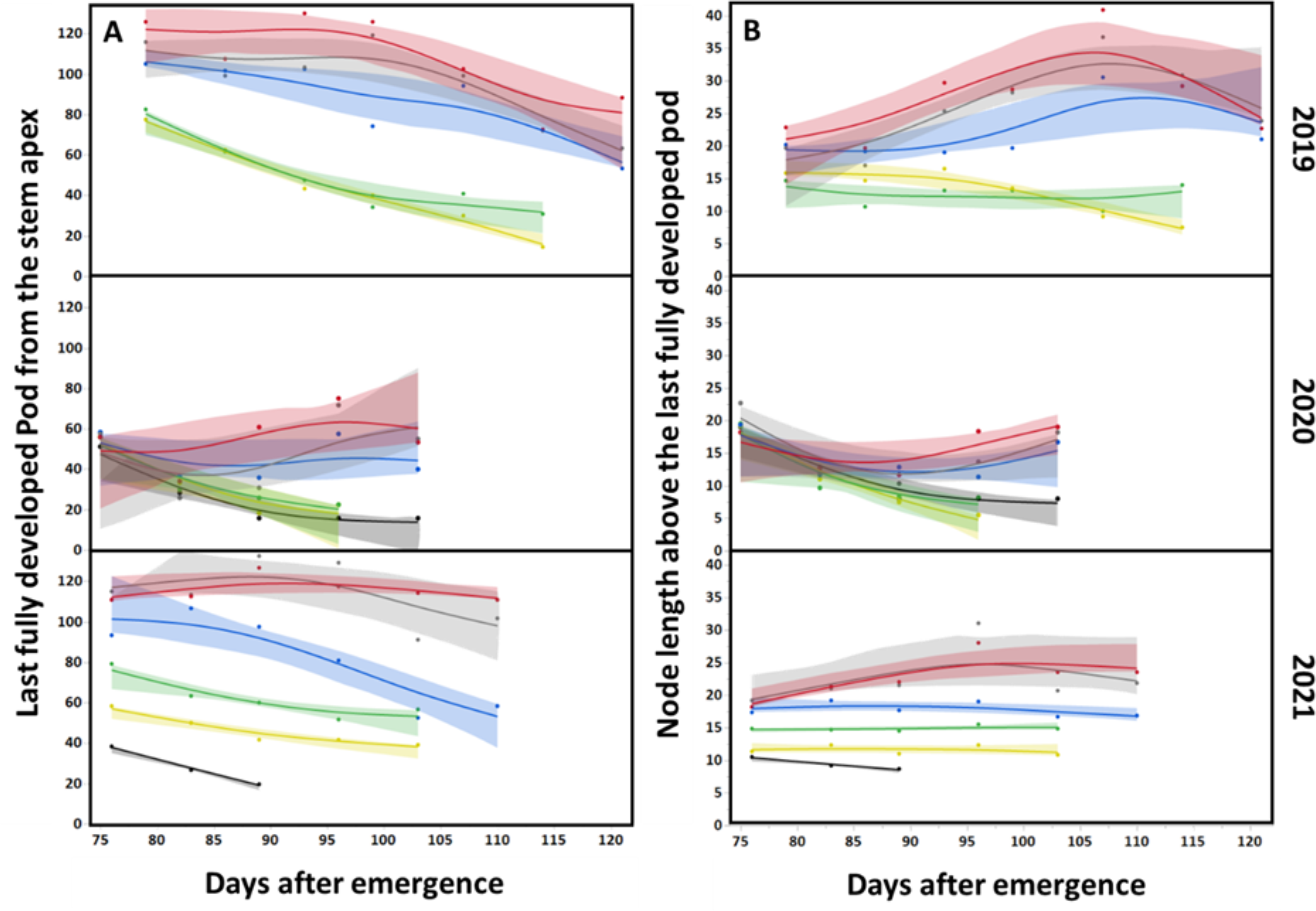
**A**) Distance of the last fully developed Pod (LFDP) from the stem apex, and **B**) node length above the last fully developed pod by irrigation treatments along the three experimental seasons (Gilat 2019 – 2021). Irrigation factor: 0 (black), 0.5 (Yellow), 0.7 (Green), 1 (blue), 1.2 (gray), and 1.4 (red). The curves represent the mean (*n*=6). Shading indicates the standard error.

The dynamic of the distance from the youngest fully inflated pod (LFDP-apex distance) to the stem apex over the crop reproductive periods are presented in Figure 3B. While all irrigation treatments start with a similar distance of LFDP-Top (around 70 mm), the irrigation treatments with IF<1 responded with a shortening of LFDP-Top values, dropping to 10-20 mm by the end of the season. Irrigation treatment with IF>1 maintained the initial LFDP-Top value and even increased it slightly by the mid-season (70-100 mm). The IF=1 treatment showed a constant decrease with a constant slope throughout the irrigation period (Fig. 3B).

### 3.3 Longitudinal pattern of dry matter accumulation

The longitudinal pattern of total dry matter (DM, manual harvested) accumulation during the three growing seasons, from about four weeks before irrigation onset until full crop maturity, is presented in Figure 4A. The seasonal accumulation of total DM differed between the irrigation treatments with the greater the amount of irrigation, the higher the DM accumulation. The seasonal patterns of the reproductive DM showed that in all treatments and during all seasons, there is a phase of about three weeks, starting from the formation of the first pods, in which pods are developing but with no appreciable reproductive DM accumulation (Fig. 4B). About two weeks after irrigation onset, this trend changes and each treatment enter a phase of rapid reproductive DM accumulation. While the water deficit treatments (IF<1) result in a shorter crop cycle and stop accumulating reproductive DM, the well-irrigated treatments (IF≥1) continue growing and set more pods (and DM) for extended periods, as reflected in the respective DM dynamics (Fig. 4B). At the end of the experiment, the plots were mechanically harvested. Generally, with the increased water availability, the grain yield increased until IF=1.2 (Fig. 4C, Table S4). The IF=1.4 treatment showed an inconsistent trend (either decrease, equally, or increased grain yield for the 2019, 2020, and 2021 seasons) compared with the IF=1.2 treatment. Notably, these differences were below the significance threshold for all seasons. The extreme weather conditions during the 2020 season (i.e., heat waves) resulted in two groups between the irrigation treatments, with IF<1 and IF≥1 clustering together. Notably, the non-irrigated treatment produced significantly lower yields in both the 2020 and 2021 seasons.

**Figure 4.**
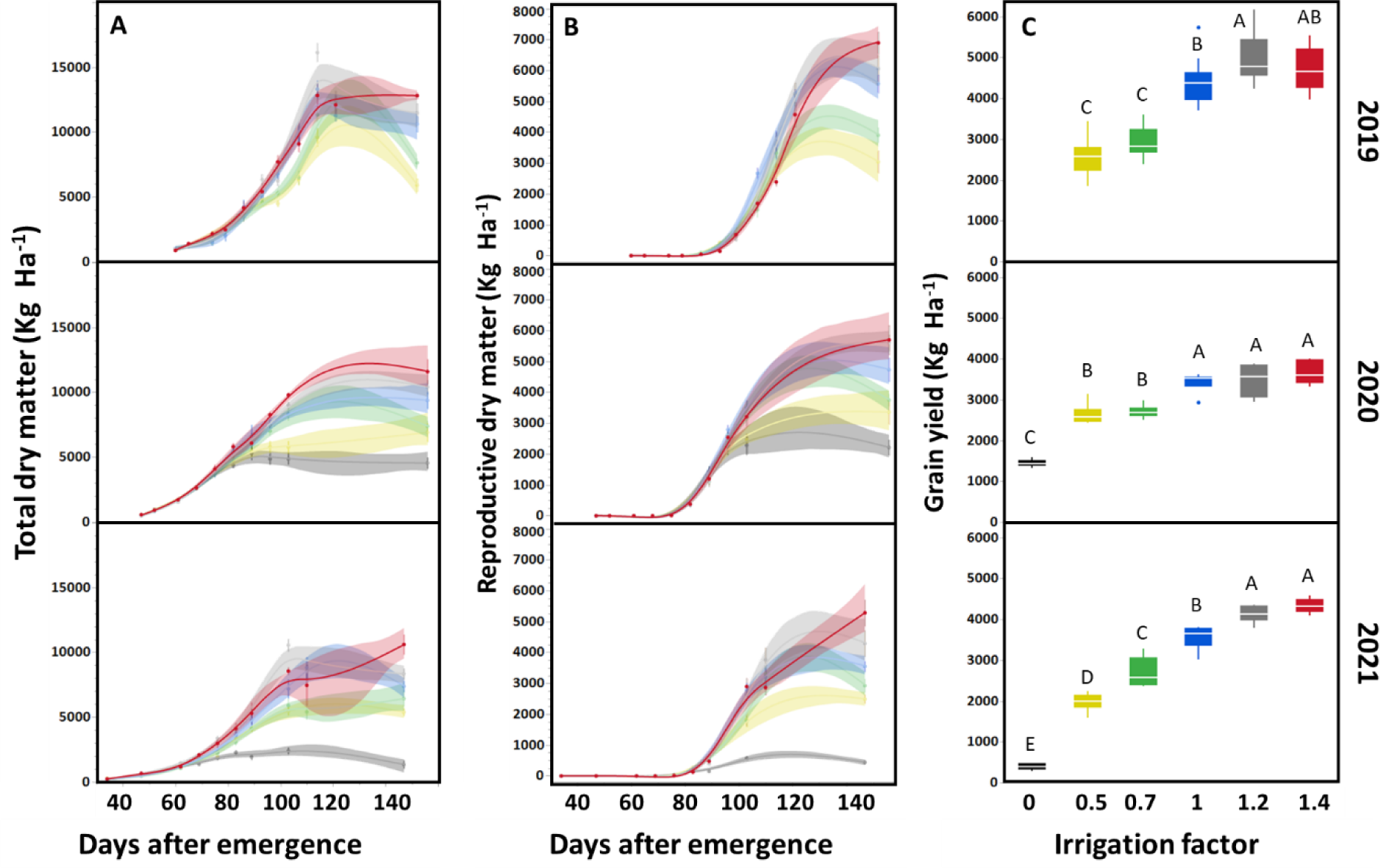
Seasonal accumulation of **A**) above-ground total dry matter (Tot. DM), **B**) reproductive DM (Rep. DM) and **C**) grain yield (GY). by irrigation treatment along the three growing seasons (Gilat 2019-2021). The curves represent the mean (*n*=6). Shading indicates the standard error. Irrigation factor: 0 (black), 0.5 (Yellow), 0.7 (Green), 1 (blue), 1.2 (gray), and 1.4 (red). Different letters indicate significant differences at *P*≤0.05.

### 3.4 Contribution of irrigation to grain yield

To test the grain yield contribution of additional water availability to grain yield, we plotted the GY along the three growing seasons based on the irrigation factor IF. While under rainfed conditions (IF=0), yield is determined solely by climatic conditions (precipitation and ET_0_ during grain filling period), under increasing irrigation treatments, we found linear responsiveness to added water across all years up to IF≤1.2 (Fig. 5). The yield differences between years are the result of environmental conditions and especially heat wave events, their timing during the irrigation period and intensity. Under irrigation treatment of IF=1.4, we found an inconsistent response that varied between years. In 2019, yield was lower than the irrigation treatment of IF=1.2, whereas, in the 2020 and 2021 seasons, the additional water contributed to increased yield gain, but with different trends (slope of graph) and with a non-significant difference from irrigation treatment IF=1.2.

**Figure 5.**
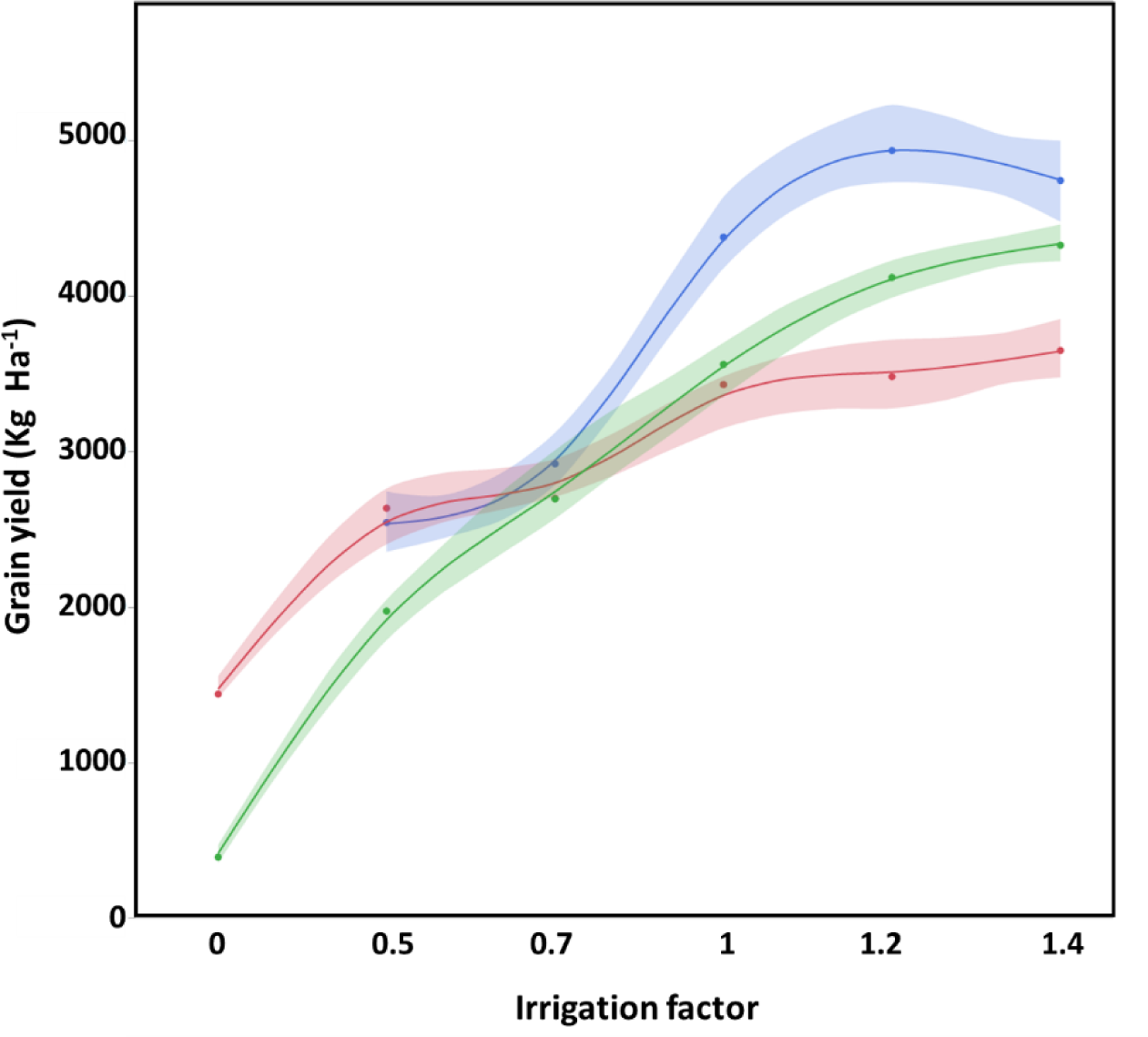
Grain yield (mechanical harvest) by irrigation treatment during the three growing seasons (2019, 2020, and 2021 (blue, red, and green, respectively) at Gilat). The curves represent the mean (*n*=6). Shading indicates the standard error.

To test the response of chickpea crop to water availability during the growing season (from sowing to maturity), we calculated for each treatment (IF 0-1.4) in each growing season (2019-2021) the available amount throughout the season (i.e., rain and irrigation applied). The intersection point (X-axis=177 mm; Fig. 6A) implies the minimal amount of water that allows the plant’s development but limits the plant’s ability to produce grains. The second parameter is the grain yield values (Y-Axix) under optimal growing conditions, as represented by the non-responsiveness of grain yield to the irrigation zone (X-axis values >580) for Israeli chickpea varieties. In semi-arid Mediterranean climate conditions, the value ranges around 4500 kg Ha^-1^. The third parameter is the slope of the graph (R^2^=0.81; Y= -1498 + 10.56X) in its linear range (water consumption 0-580 mm), which expresses the water productivity under specific climate and growing conditions (Fig. 6A).

**Figure 6.**
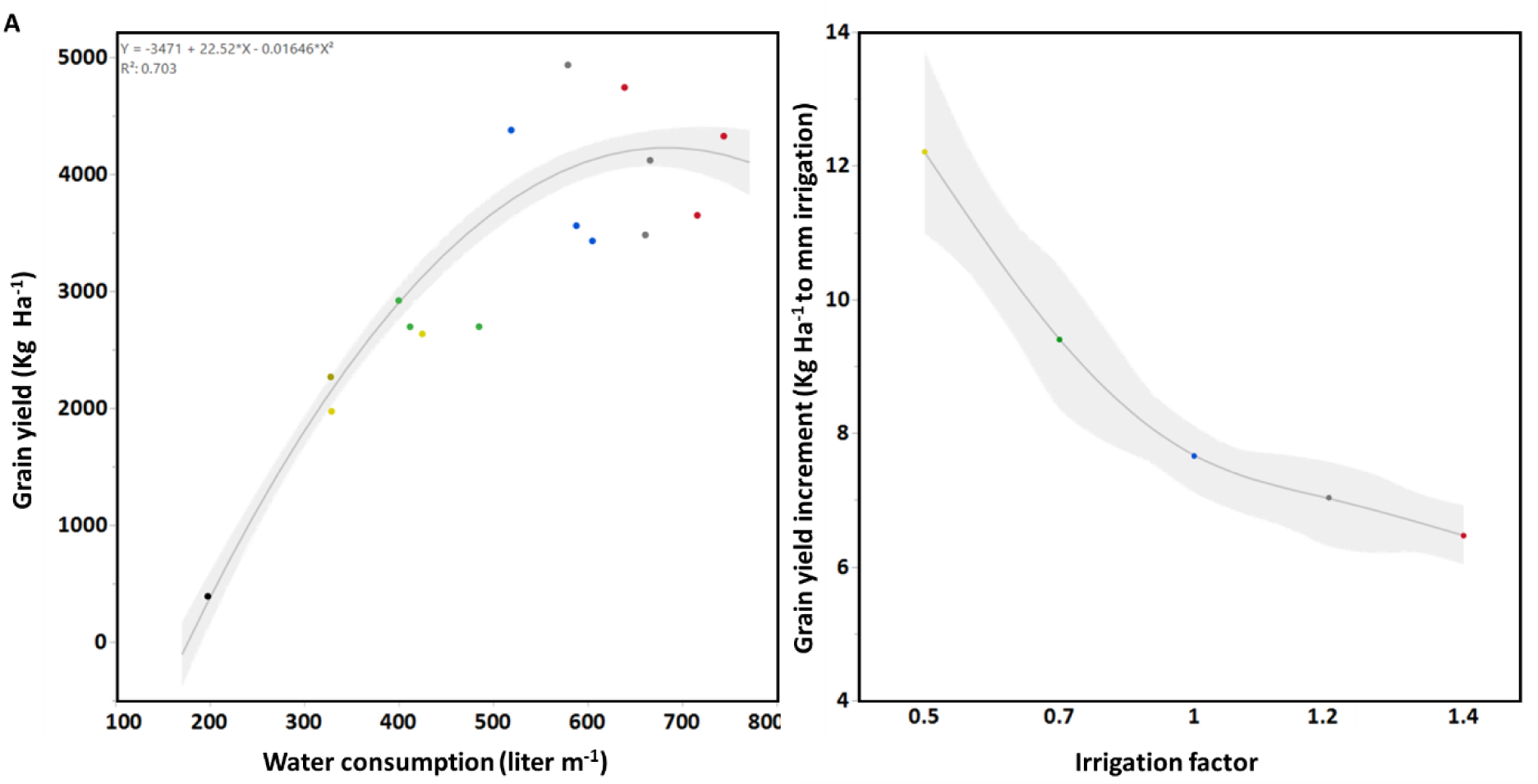
**A**) Grain yield (mechanical harvest) response to water consumption over 2019, 2020, and 2021 growing seasons at Gilat, Israel. **B**) Grain yield (average of 2020 and 2021 growing seasons) contribution to mm (liter m^-1^) irrigation by irrigation factor (IF). The curves represent the mean (*n*=6). Shading indicates the standard error.

The contribution of additional water to grain yield accumulation under each irrigation treatment (GY_Ifx_ – GY_IF0_) / mm of irrigation_Ifx_) is presented in Figure 6B. Under the lowest water availability (i.e., lower IF), the highest contribution to yield was 8-12 kg grain ha^-1^ mm^-1^ of irrigation. With increasing water availability (IF≥1), the contribution to yield per mm irrigation water decreased to 6-8 kg grain ha^-1^ mm^-1^. However, it still allows additional yield accumulation and offers an approach for achieving the maximum grain yield potential of the field.

## 4. Discussion

The unpredictable and fluctuating climatic conditions associated with present and projected climate change have become the primary constraint for chickpea production worldwide, jeopardizing future food security. This study’s primary objective was to develop an optimal strategy for chickpea irrigation based on meteorological data and supported by morpho-physiological parameters to promote maximizing grain yields under the semi-arid Mediterranean basin. We have used the measured ET_c_ values to design differential irrigation regimes well reflected in differential crop physiology and morphological parameters (Tables 1, S2; Figs. 2-3). Eventually, these irrigation treatments resulted in a range of biomass and grain yield values (Fig. 4), which may assist in formulating chickpea drip-irrigation protocols for semi-arid Mediterranean growing regions.

### 4.1. Morphological- rather than physiological- traits easily capture the crop water status dynamics

Assessing the crop water status during the growing season is expensive, cumbersome, and time-consuming, which poses a challenge for plant physiologists and farmers. Various strategies have been suggested, including direct sensors (e.g., soil tensiometers, stem diameters, pressure chamber, etc.) or remote sensing (e.g., multi-spectral and thermal imaging) to assess this trait (Zhou et al., 2021; Ezenne et al., 2019; Avneri et al., 2023). In the current study, we used a pressure chamber to measure directly the crop leaves water potential in the field (Fig. 2). This strategy was used to understand the plant-water relationships (Turner, 1988). However, this technology does not enable many measurements during any sampling day. The need for real-time knowledge of the crop water status emphasizes the necessity to identify objective morpho-physiological indices to determine the plant water status (Cohen et al., 2015). Our irrigation protocol and the documented morpho-developmental indices of node length above the last fully developed pod (LFDP) and distance between LFDP to plant apex can help by providing “morphological” feedback to the irrigation schedule that enables monitoring the crop water status without sophisticated equipment. When considering the overall results of node length above LFDP and distance LFDP-plant apex between years and irrigation treatments, it seems that node length shorter than 20 mm indicates that the plant experiences water stress. In comparison, LFDP-plant apex values above 30 mm may hint at excess irrigation.

Under Mediterranean-basin climatic conditions for Israeli Kabuli-type varieties, node length above LFDP around 25 mm is required to maximize grain yield. Higher values of our morpho-developmental parameters may serve as an indication to decrease the amount of irrigation in the following application. In comparison, lower values indicate water stress and higher irrigation cubature should be considered. In addition, a farmer may monitor these parameters on his chickpea crop over several seasons and varieties and, if needed, make required local adjustments. In our view, the morphological indices best reflected the crop water status since plant development in effect (and by definition) integrates all environmental determinants over the growing period. Therefore, monitoring the ET_0_ and growth parameters may assist in determining the recommended water dose before each irrigation pulse, thereby ensuring uninterrupted yield accumulation and preventing yield losses from lodging, pod shading, and/or leaf diseases.

The distance between the LFDP to the plant apex reflects, in our opinion, the long-term water status throughout the podding period. This parameter shows a similar trend between years in which distance shortens as the water availability to the plant decreases and as irrigation season approaches its end (Fig. 3). The values of this parameter were statistically different between irrigation treatments in all growing seasons (Table S2), thereby providing an essential and easy measure to assess the crop water status. While irrigation treatments with IF≥1 can maintain this distance (even for 1-2 weeks) or even increase it, irrigation treatments with IF<1 are characterized by decreasing values of this parameter. This fact implies that the crop water requirements during podding are higher than the ET_0_ (IF=1), and the field should be irrigated above this value to extend the irrigation period as long as the crop can add fertile pods. Likewise, the node length above LFDP is also a reliable indicator of plant water status. This is the last stem node that ended its elongation and reflects the plant’s water availability during the period between node formation and measurement day-final length. Unlike pressure chamber measurements that provide instantaneous (day-time) information, this node length may be considered as an integrator of the conditions experienced by the plant ever since its formation to the day of measurement. No less important, it is easy to measure, requires no expensive equipment, and is not affected by extreme climate conditions that may occur on the measurement day or hour. However, it is necessary to consider that desired values may change between cultivars, growth conditions, and their interactions. This parameter is proportional to the applied water cubature, with the highest amount of irrigation, the longest the nodes. From the irrigation onset date, values are decreasing under irrigation treatments with IF <1 while irrigation treatments with IF>1 show increased length to desired values (above 20 mm). This corroborates our claim that the irrigation water demand by the crop during the podding phase (mid-season) is greater than IF>1.

Our results indicate a close match between the seasonal pattern of the two morphometric parameters (as an indication of the crop water status) and the reproductive DM accumulation across irrigation treatments and years (Fig. 4). The reliability of our morphological parameters and their usefulness in general irrigation design (and ad-hoc decision making) is evident from the close match between the dynamics of the seasonal leaf water potential values and the longitudinal patterns of the LFDP and the length of the node above the LFDP.

### 4.2 Extentending the reproductive phase is essential for grain yield improvement under semi-arid conditions

Oweis et al. (2004) suggested that 2-3 additional irrigations per season during the reproductive phase enable higher chickpea grain yield in the Mediterranean basin. We have used a different approach of more frequent irrigations to maintain a constant soil moisture content to stabilize the crop water status for extended periods. We suggest that to achieve high grain yields (>4 Ton Ha^-1^), the irrigation period must be extended by keeping the crop vital and functioning. Davies et al. (1999) showed a correlation between leaf water potential and net photosynthesis (Davies et al., 1999). Thus, by maintaining a desirable plant’s water status, the plant can support a high level of net photosynthesis that contributes to appropriate partitioning and redistribution of dry matter from vegetative organs (stems and leaves) to reproductive organs (flowers and pods). Indeed, this is one of the main physiological determinants of high grain yield in chickpeas growing under Mediterranean-type conditions (Leport et al., 1999). Extending reproductive growth in crops with indeterminate growth patterns (Bonfil and Pinthus, 1995), which support high grain yield, can only be achieved by carefully tuning irrigations at high frequency. Such an approach may enable farmers to match crops’ water demand via quick response during extreme weather conditions. Adequate water status allows evaporative cooling of the foliage during extreme weather conditions, thereby keeping plants active and functional. Indeed, our IF=1.2 and 1.4 treatments were capable of preventing crop collapse and growth cessation during heat waves with days above 40°C (Fig. S1). In other words, maintaining a long crop growth period will likely result in large and well-developed plants that can bear more main and secondary branches carrying flowers and pods (Pushpavalli et al., 2015).

### 4.3 IF=1.2 provides the most productive regime for chickpea in Mediterranean conditions

The contribution of the amount of irrigation to grain yield was defined only in 2020 and 2021 (with rainfed treatment) by calculating the yield differences between specific irrigation treatment and the rainfed treatment, divided by the amount of supplied water (Fig. 6A). Notably, under low IF (e.g., 0.5 and 0.7) irrigation treatments, additional water had a relatively large contribution to the grain yield (∼10 kg grains Ha^-1^ mm^-1^ irrigation water). Under high IF≥1 irrigation levels (1, 1.2, and 1.4), water contribution to yield decreased to about 7.0 kg grains Ha^-1^ mm^-1^ irrigation. We suggest that using an irrigation factor of ca. 1.1-1.2 may support relatively high grain yield levels (>4 t/ha) even in years with high intensity of extreme heat waves, such as the 2020 season (Fig. S1), albeit at the cost of low water-use-efficiency.

The current chickpea irrigation protocol (Allen et al., 1998) is based on ET_0_ multiplied by the crop coefficient Kc=1 during the mid-season irrigation period and 0.35 during the late-season irrigations to supply the crop water demand. Rinaldi et al. (2008) suggested an IF of 0.97 over mid-season and IF of 0.54 at late-season for Desi-type chickpea. On the other hand, our findings suggest that GY could be maximized by implementing an IF of 1.2 from the onset of irrigation until the end of the growing season. Parallel monitoring of the morphological parameters offers feedback for fine-tuning the irrigation’s cubature. A possible explanation for our high IF compared with previous studies (Rinaldi et al. 2008; Allen et al., 1998) is probably due to the harsh weather conditions (high temperatures, ET_0_ and VPD values) that prevailed during our growing seasons, which shift the ETc to higher levels as described in Figure 21 (Allen et al., 1998). Additional research is needed to evaluate the different types of cultivars (Kabuli *vs*. Desi) yield accumulation and morphological values response to different irrigation factors.

### 4.4 Conclusion and future perspective

Extending the reproductive period from the beginning of the spring to early summer is essential for producing high yields in the Mediterranean basin (i.e., rising temperatures and extremely hot days) (Fig. S3A). The intensity and duration of heat waves can result in dramatic yield reduction due to the shortening of the reproductive period, as happened in the 2020 season (Fig. 4). Such high-evaporative requirement days cause flowers and young pods abortion and decrease the plant’s turgor, which may cause crop lodging (see Fig. S3B). This phenomenon is irreversible, and chickpea plants usually cannot resurrect, making the continuation of irrigation ineffective. Thus, maintaining the crop fully turgid throughout the irrigation period may enable plants to cope with extreme heat waves. The morphological traits are a powerful feedback tool for examining the applied irrigation water and allow a fine adjustment under changing growing conditions such as crop rotation, field patchiness, and different locations. The morphological traits reflect the effects of a two weeks period, unlike single time-point physiological characteristics, and therefore offer a more reliable tool for plant water-status evaluation.

Our new irrigation guidelines for semi-arid environments for chickpea were already adopted by farmers in commercial fields (including for other kabuli cvs. than Zehavit) in southern Israel (2021 and 2022 rainfall 200-450 mm) during the 2021 and 2022 growing seasons, resulting in high yields ranging from 4.0-5.5 Ton ha^-1^ (Fig. S3C). Notably, this irrigation practice efficiently coped with extremely high daily temperatures (> 40°C) during the irrigation periods, which is expected to happen more often in the Mediterranean basin. While this protocol resulted in increased yields, it is important to point out that application of our guidelines to other cultivars and environmental conditions may require adjusting the use of the morphological parameters. Moreover, this experimental approach of differential irrigation method, while each irrigation treatment receives the desired amount of water accurately for the same irrigation duration, could also be implemented for other crops for a better understanding of their irrigation requirements. Importantly, our findings and the increased frequency of extreme weather events call for an urgent re-evaluation of current irrigation factors and protocols to cope with the current and projected climatic conditions and enhance yields to ensure food security.

## Abbreviations

DAE: Days after emergence
ET: Evapotranspiration
g: Gram
Ha: Hectare
IF: Irrigation factor
kg: Kilogram
LFDP: Last Fully Developed Pod
Ψ^LWP^: Leaf water potential
max: Maximum
mm: Millimeter
min: Minimum
ETo: Reference crop evapotranspiration
temp: Temperature
t: Ton
WP: Water productivity

## Acknowledgments

We thank Prosperi Abonete, Gabriel Mulero, Yuval Siboni, and Yehudit Shamir for assisting with the field experiments. This research was supported by the State of Israel Ministry of Agriculture and Rural Development (grant # 20-02-0087). Shahal Abbo is the incumbent of The Jacob & Rachel Liss Chair in Agronomy, the Hebrew University of Jerusalem.

