## Supplementary material for "Optimization of chickpea irrigation in a semi-arid climate based on morpho-physiological parameters": SI data

### Supplementary materials

**Table S1.** Summary of climatic and irrigation data during the chickpea growth season by year (2019-2021), Gilat, Israel. Average minimum (min.) and maximum (max.) temperatures (temp.) through the growing season, until irrigation onset and during the irrigation period. Amount of precipitation, sprinklers, and drip irrigation (Irrigation factor, IF=1). Total water application (precipitation and irrigation). Evapotranspiration from emergence to maturity and from irrigation onset to maturity.

| | **2019** | **2020** | **2021** |
|---|---|---|---|
| Average min. temp. ($^oC$) | 11.30 | 11.30 | 11.30 |
| Average max. temp. ($^oC$) | 25.70 | 26.10 | 26.00 |
| Average min. temp. until irrigation onset ($^oC$) | 9.36 | 10.14 | 8.74 |
| Average max. temp. until irrigation onset ($^oC$) | 21.75 | 23.44 | 21.83 |
| Average min. temp. irrigation period ($^oC$) | 14.70 | 14.38 | 14.20 |
| Average max. temp. irrigation period ($^oC$) | 32.84 | 33.00 | 30.88 |
| Precipitation (mm) | 130.2 | 225.6 | 140.7 |
| Sprinkler irrigation (mm) | 85 | 25 | 60 |
| Dripping irrigation IF=1 (mm) | 302 | 277.5 | 389.5 |
| Total water application (mm) | 517.2 | 528.1 | 590.2 |
| Irrigation period (days) | 42 | 28 | 54 |
| Evapotranspiration from emergence to maturity (mm) | 574 | 478.8 | 561.9 |
| Evapotranspiration from irrigation onset to maturity (mm) | 310.9 | 228.7 | 373.2 |

**Table S2.** Analysis of variance (ANOVA) for total dry matter (Tot. DM), reproductive DM (Rep. DM), grain yield (GY), harvest index (HI), Leaf water potential ($\Psi^{LWP}$), water productivity (WP), the distance of the last fully developed pod (LFDP) from plant-apex and node length above LFDP at median irrigation period, under different irrigation regimes (irrigation factor, IF=0.5, 0.7, 1, 1.2, and 1.4) for each experimental season (2019, 2020, and 2021).

| Source of variation | | Tot. DM | | | | | Rep. DM | | | | | GY | | | | | $\Psi^{LWP}$ | | | | |
|---|---|---|---|---|---|---|---|---|---|---|---|---|---|---|---|---|---|---|---|---|---|
| | | d.f | 2019 | d.f | 2020 | 2021 | d.f | 2019 | d.f | 2020 | 2021 | d.f | 2019 | d.f | 2020 | 2021 | d.f | 2019 | d.f | 2020 | 2021 |
| Irrigation trearment | sum of squares | 4 | 1.94E+08 | 5 | 2.00E+08 | 2.93E+08 | 4 | 60200000 | 5 | 5.61E+07 | 8.29E+07 | 4 | 28700000 | 5 | 20600000 | 66900000 | 4 | 79.801 | 5 | 1217.16 | 643.852 |
| | Probe > F | | 0.0001 | | 0.0001 | 0.0001 | | 0.0001 | | 0.0001 | 0.0001 | | 0.0001 | | 0.0001 | 0.0001 | | 0.0001 | | 0.0001 | 0.0001 |
| Block | sum of squares | 5 | 5865197 | 5 | 1.05E+07 | 8804330 | 5 | 2231701 | 5 | 2639446 | 2663956 | 5 | 1680698 | 5 | 271102 | 481052 | 5 | 1.14315 | 5 | 9.20333 | 58.6451 |
| | Probe > F | | 0.651 n.s | | 0.546 n.s | 0.3278 n.s | | 0.7154 n.s | | 0.6028 n.s | 0.3321 n.s | | 0.0509 | | 0.5725 n.s | 0.1496 n.s | | 0.382 n.s | | 0.046 | 0.0164 |
| Error | sum of squares | 19 | 35033578 | 25 | 63713663 | 36002265 | 19 | 15410145 | 25 | 17907154 | 10981285 | 19 | 2492762 | 25 | 1734307 | 1337037 | 19 | 3.870517 | 25 | 17.26 | 80.70765 |

| Source of variation | | LFDP | | | | | Node distance above LFDP | | | | | WP | | | | | HI | | | | |
|---|---|---|---|---|---|---|---|---|---|---|---|---|---|---|---|---|---|---|---|---|---|
| | | d.f | 2019 | d.f | 2020 | 2021 | d.f | 2019 | d.f | 2020 | 2021 | d.f | 2019 | d.f | 2020 | 2021 | d.f | 2019 | d.f | 2020 | 2021 |
| Irrigation trearment | sum of squares | 4 | 44163.3 | 5 | 7981.25 | 64495 | 4 | 1373.13 | 5 | 141.889 | 899.889 | 4 | 0.11108 | 5 | 0.15287 | 0.84368 | 4 | 0.0031 | 5 | 0.01083 | 0.23581 |
| | Probe > F | | 0.0001 | | 0.0001 | 0.0001 | | 0.0001 | | 0.0002 | 0.0001 | | 0.0454 | | 0.0001 | 0.0001 | | 0.8602 n.s | | 0.9069 n.s | 0.0001 |
| Block | sum of squares | 5 | 256.667 | 5 | 89.5833 | 769.667 | 5 | 18.1667 | 5 | 3.88889 | 21.8889 | 5 | 0.00211 | 5 | 0.01173 | 0.02635 | 5 | 0.02623 | 5 | 0.02389 | 0.00186 |
| | Probe > F | | 0.295 n.s | | 0.7741 n.s | 0.5355 n.s | | 0.3193 n.s | | 0.9588 n.s | 0.3977 n.s | | 0.9075 n.s | | 0.543 n.s | 0.0942 n.s | | 0.0973 n.s | | 0.6521 n.s | 0.9853 n.s |
| Error | sum of squares | 19 | 776.667 | 25 | 897.9167 | 4593.333 | 19 | 57.6667 | 25 | 96.4444 | 101.7778 | 19 | 0.825256 | 25 | 0.071 | 0.061669 | 19 | 0.04813 | 25 | 0.179132 | 0.071121 |

**Table S3. Data of morphological traits:** the distance of the last fully developed pod (LFDP) from plant-apex and node length above LFDP at median irrigation period**.** Data is mean (*n*=6)±SE.

| **Distance of the last fully developed pod from plant-apex (mm)** | | | | | | | |
|---|---|---|---|---|---|---|---|
| **2019 season** | | | | | | | |
| | **79** | **86** | **93** | **99** | **107** | **114** | **121** |
| **IF=0.5** | 77.5±4.3 | 62.5±3.4 | 43.3±2.5 | 40.0±2.2 | 30.0±3.2 | 14.5±0.5 | NA |
| **IF=0.7** | 82.5±4.6 | 61.7±3.3 | 47.5±2.1 | 34.2±2.4 | 40.8±3.0 | 30.8±1.5 | NA |
| **IF=1** | 105.0±4.1 | 101.8±5.3 | 102.5±3.4 | 74.2±1.5 | 94.2±2.4 | 72.5±1.7 | 53.3±4.4 |
| **IF=1.2** | 115.8±4.7 | 99.2±9.1 | 103.3±2.1 | 119.2±3.0 | 99.2±1.5 | 72.0±2.4 | 63.3±5.6 |
| **IF=1.4** | 125.8±8.7 | 107.5±5.3 | 130.0±3.4 | 125.8±3.5 | 102.5±3.1 | 72.5±1.1 | 88.3±4.9 |
| **Season 2020** | | | | | | | |
| | **75** | **82** | **89** | **96** | **103** | | |
| **IF=0** | 51.2±2.4 | 28.3±2.5 | 15.8±0.8 | NA | NA | | |
| **IF=0.5** | 56.2±3.8 | 33.3±5.1 | 18.3±1.05 | 22.5±1.7 | NA | | |
| **IF=0.7** | 55.8±0.8 | 30.0±2.2 | 25.8±1.5 | 22.5±1.1 | NA | | |
| **IF=1** | 58.3±2.5 | 36.7±4.4 | 35.8±3.7 | 57.5±1.7 | 40.0±2.9 | | |
| **IF=1.2** | 57.5±2.8 | 25.8±1.5 | 30.8±2.0 | 71.7±1.0 | 55.0±5.2 | | |
| **IF=1.4** | 55.8±3.0 | 34.2±4.5 | 60.8±3.3 | 75.0±1.8 | 53.3±6.7 | | |
| **Season 2021** | | | | | | | |
| | **76** | **83** | **89** | **96** | **103** | **110** | |
| **IF=0** | 38.3±2.1 | 26.7±2.8 | 19.7±2.7 | NA | NA | NA | |
| **IF=0.5** | 58.3±5.7 | 50.0±4.6 | 41.7±6.0 | 41.7±1.7 | 39.2±5.2 | NA | |
| **IF=0.7** | 79.2±4.4 | 63.3±7.7 | 60.0±2.2 | 51.7±3.1 | 56.7±6.7 | NA | |
| **IF=1** | 93.3±4.9 | 106.7±6.5 | 97.5±3.3 | 80.8±5.1 | 52.5±2.81365 | 58.3±7.5 | |
| **IF=1.2** | 115.0±2.6 | 113.3±7.9 | 132.5±10.1 | 129.2±9.6 | 91.1±17.2 | 101.7±3.1 | |
| **IF=1.4** | 110.8±7.0 | 112.5±6.5 | 126.7±4.0 | 117.5±6.7 | 114.2±6.6 | 110.8±4.2 | |

| Node length above LFDP (mm) | | | | | | | |
|---|---|---|---|---|---|---|---|
| **2019 season** | | | | | | | |
| | **79** | **86** | **93** | **99** | **107** | **114** | **121** |
| **IF=0.5** | 15.8±0.9 | 14.7±1.2 | 16.5±0.7 | 13.5±1.0 | 9.2±0.5 | 7.5±0.5 | NA |
| **IF=0.7** | 14.7±0.8 | 10.7±1.5 | 13.2±0.6 | 13.2±0.9 | 10.0±0.0 | 14.0±0.6 | NA |
| **IF=1** | 20.2±1.1 | 19.2±0.5 | 19.0±0.4 | 19.7±0.3 | 30.5±1.6 | 29.2±1.5 | 21.0±0.4 |
| **IF=1.2** | 19.7±0.8 | 17.0±1.5 | 25.3±1.3 | 28.2±0.7 | 36.7±1.1 | 30.8±0.8 | 23.8±1.8 |
| **IF=1.4** | 22.8±1.1 | 19.7±0.3 | 29.7±0.6 | 28.7±0.4 | 40.8±2.0 | 29.2±0.8 | 22.7±1.1 |
| **Season 2020** | | | | | | | |
| | **75** | **82** | **89** | **96** | **103** | | |
| **IF=0** | 19±1.2 | 12.7±1.5 | 8.0±0.4 | NA | NA | | |
| **IF=0.5** | 19.2±1.2 | 11.0±1.3 | 7.5±0.7 | 5.5±0.3 | NA | | |
| **IF=0.7** | 19.5±0.9 | 9.7±0.3 | 8.3±0.8 | 8.2±0.4 | NA | | |
| **IF=1** | 19.3±1.4 | 11.7±0.1 | 12.8±1.1 | 11.3±0.4 | 16.7±0.8 | | |
| **IF=1.2** | 22.7±1.1 | 12.0±1.0 | 10.3±0.3 | 13.7±0.9 | 18.2±1.2 | | |
| **IF=1.4** | 18.2±0.6 | 12.8±1.01 | 11.7±0.8 | 18.3±1.0 | 19±1.1 | | |
| **Season 2021** | | | | | | | |
| | **76** | **83** | **89** | **96** | **103** | **110** | |
| **IF=0** | 10.5±0.8 | 9.2±1.1 | 8.7±0.7 | NA | NA | NA | |
| **IF=0.5** | 11.3±1.2 | 12.3±0.8 | 11.0±0.9 | 12.3±0.9 | 10.8±1.2 | NA | |
| **IF=0.7** | 14.8±1.1 | 14.7±1.2 | 14.5±0.5 | 15.5±0.7 | 14.8±0.8 | NA | |
| **IF=1** | 17.3±0.9 | 19.2±0.4 | 17.7±0.6 | 19.0±1.2 | 16.7±0.6 | 16.8±0.7 | |
| **IF=1.2** | 19.2±1.2 | 21.0±0.5 | 21.5±0.8 | 31.0±2.0 | 20.7±0.7 | 21.8±0.8 | |
| **IF=1.4** | 18.2±1.1 | 21.3±1.2 | 22.0±1.2 | 28.0±0.8 | 23.5±0.4 | 23.5±1.4 | |

**Table S4:** Average chickpea grain yields kg $Ha^{-1}$ (mechanically harvested) by irrigation factor (IF), during the 2019, 2020, and 2021 growing seasons at Gilat, Israel.

| Irrigation factor | Growing season | | |
|---|---|---|---|
| | 2019 | 2020 | 2021 |
| **IF=1** | NA | 1439.7 C | 390.7 E |
| **IF=0.5** | 2544.0 C | 2635.9 B | 1973.3 D |
| **IF=0.7** | 2921.5 C | 2697.8 B | 2696.7 C |
| **IF=1** | 4379.2 B | 3431.4 A | 3561.3 B |
| **IF=1.2** | 4936.6 A | 3482.3 A | 4120.7 A |
| **IF=1.4** | 4744.1 AB | 3650.6 A | 4327.5 A |

Different letters indicate a statistical difference between irrigation treatment by Tukey test at *P*=0.05.

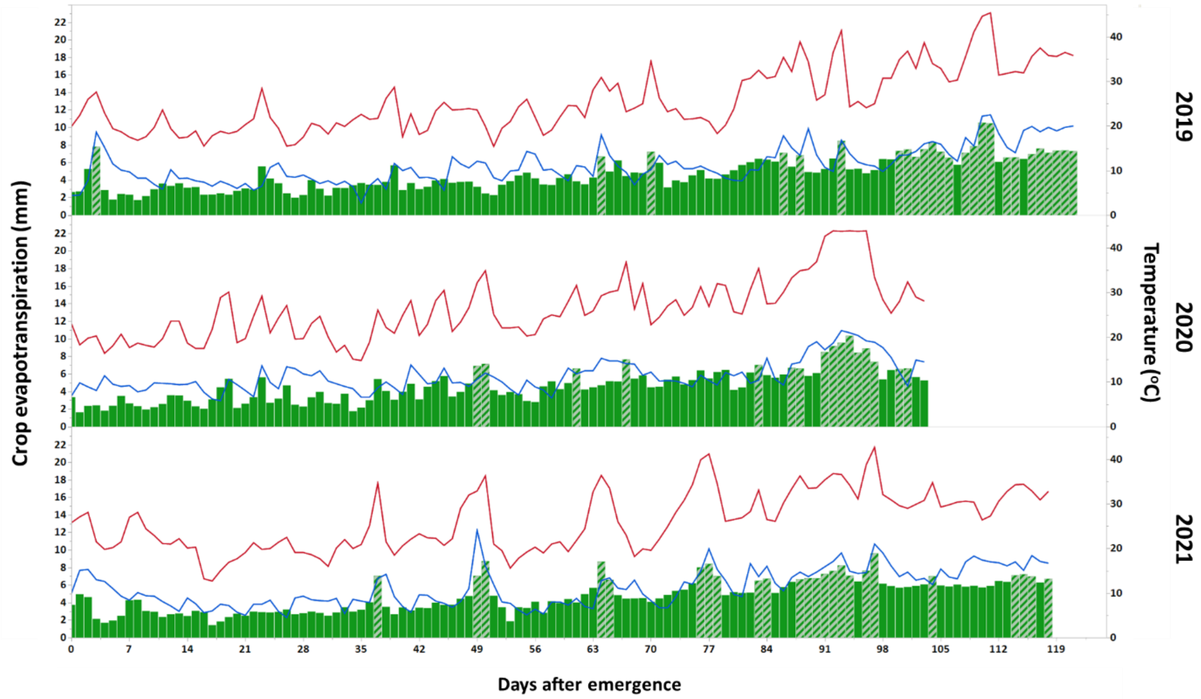


**Figure S1.** Seasonal pattern of daily minimum (blue) and maximum (red) temperatures and a histogram of the daily crop evapotranspiration (ETc; green) during the 2019-2021 chickpea growth season, Gilat, Israel. Days with ETc>6.5mm (i.e., heat-wave) are marked.

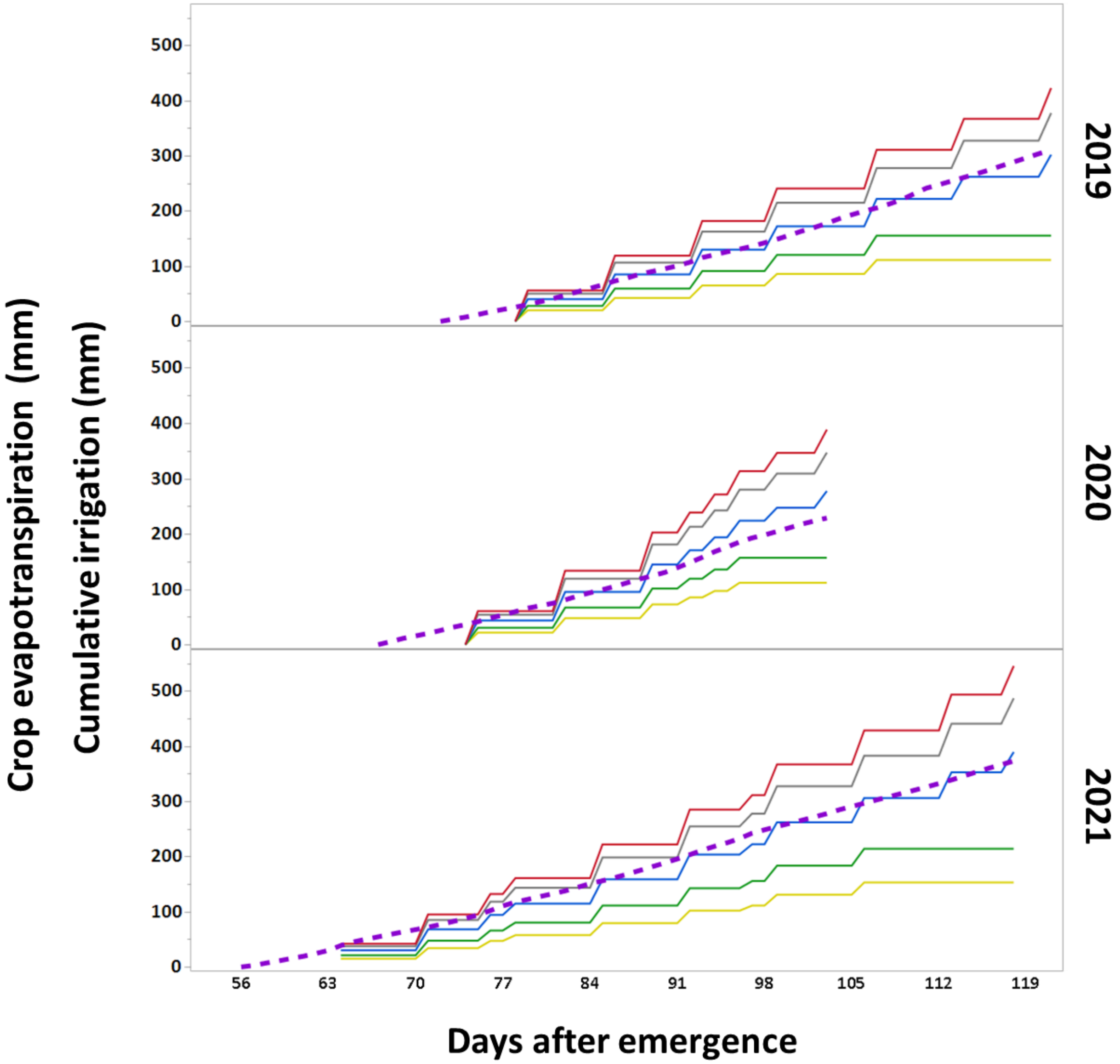


**Figure S2.** Seasonal cumulative evapotranspiration (purple) and irrigation by treatments, starting one week before irrigation onset until the last irrigation, Gilat (2019-2021). Irrigation factors 0 (black), 0.5 (Yellow), 0.7 (Green), 1 (blue), 1.2 (gray) and 1.4 (red).

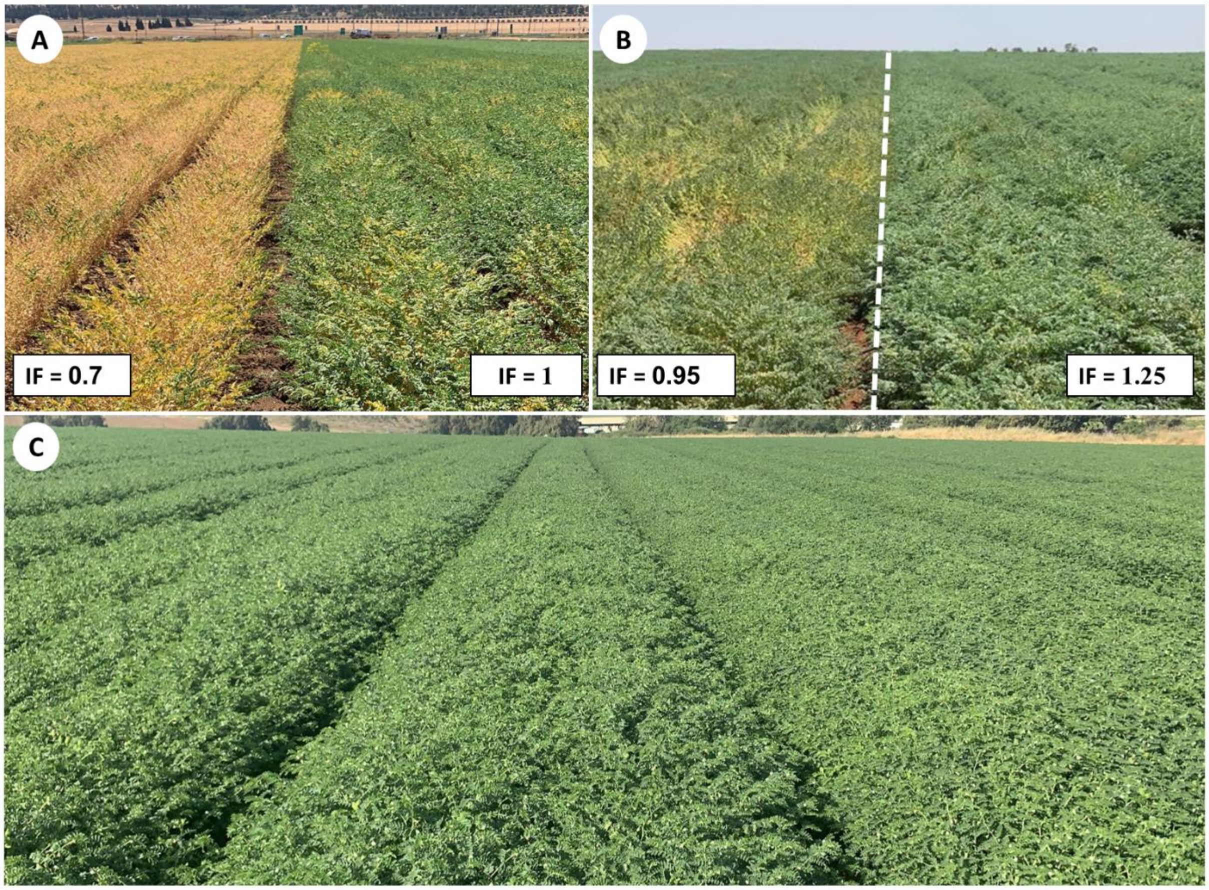


**Fig. S3.** **A**) Commercial field *cv*. Yarden (Beit Keshet, northern Israel) during the 2021 growing season. Two irrigation factors: IF=0.7 *vs*. IF=1. **B**) Commercial field *cv*. Zehavit (Geva, northern Israel) during the 2021 growing season. Two irrigation factors: IF=0.95 *vs*. IF=1.25. **C**) Commercial field *cv*. Zehavit (Or Ha'nner, southern Israel) during the 2021 growing season, with optimal irrigation (IF=1.25). The average yield at this location was >4.8 tons $Ha^{-1}$.
